# Assessing the impact of mono- and bi-allelic deletions in *NRXN1* on synaptic function

**DOI:** 10.64898/2026.02.20.707024

**Authors:** Aicha Massrali, Arkoprovo Paul, Rugile Matuleviciute, Nicholas JF Gatford, Lucia Dutan-Polit, Shekhar Kedia, Saifur Rahman, Deepak P. Srivastava, Mark Kotter, Deep Adhya, Simon Baron-Cohen

## Abstract

Neurexin 1 (NRXN1) is an adhesion protein involved in synapse development and function. Mutations in *NRXN1* are strongly linked with neurodevelopmental and psychiatric conditions. Mono-allelic *NRXN1* mutations are associated with autistic traits, with increased likelihood of co-occurring intellectual disability. However, mono-allelic mutations have variable penetrance and occur in individuals without neurodevelopmental phenotypes. Conversely, bi-allelic mutations, though rarer, are associated with more stable penetrance and severe neurodevelopmental phenotypes. Human induced pluripotent stem cells (iPSC) have been used to study how mutations in *NRXN1* impacts its function, with most studies focusing on monoallelic mutations. In this study, we systematically compared monoallelic and biallelic mutations in *NRXN1,* characterising their effects on molecular, synaptic, and functional phenotypes. Using CRISPR-Cas9, we introduced indels in *NRXN1* exon 19, in an iPSC line containing inducible *NGN2*. These edits caused either mono-allelic or compound bi-allelic frameshift mutations. iPSCs containing either mutation robustly generated glutamatergic neurons, but these neurons displayed reduced expression of major *NRXN1* isoforms. Transcriptomic profiling revealed modest gene expression changes in mono-allelic mutant neurons, whereas bi-allelic mutants exhibited extensive dysregulation of gene networks associated with neuronal maturation and synaptic function. Furthermore, synaptic phenotypes were mild in mono-allelic mutants but pronounced in bi-allelic mutant neurons. Both mono-allelic and bi-allelic mutant neurons displayed alterations in neuronal network activity and reduced peak depolarisation responses to KCl stimulation. Together, these data demonstrate that *NRXN1* exhibits gene-dosage sensitivity, with bi-allelic disruption of exon 19 unmasking molecular, synaptic, and functional phenotypes that are only modest in mono-allelic mutant neurons.

## Introduction

Neurexins, including Neurexin 1 (NRXN1), are enriched at pre-synaptic terminals, where they function as pre-synaptic adhesion molecules with key roles in synapse assembly and function ^1,2^. Neurexins form calcium-dependent complexes with a diverse repertoire of binding partners including neuroligins. It is through these interactions that NRXN1 plays a key role in synapse formation, organisation of synapses and maintaining structural stability ^1,3^. Genetic variation in the *NRXN1* gene, located on chromosome 2p16.3, is strongly associated with a range of neurodevelopmental and psychiatric conditions, including autism, schizophrenia and intellectual disability (ID) ^1,4^. *NRXN1* has an Eagle (Evaluation of Autism Gene Link Evidence) score of 143.75 and is thought to play a role in autism aetiology. It is estimated that 24% of autistic individuals with *NRXN1* mutations have ID, although this may be underestimated as 62% of the NRXN1-autistic population were unable to complete a test for ID ^2^.

Evaluation of *NRXN1* mutations and associated neurological conditions revealed deletions in exons 6-24 had the greatest association with intellectual disabilities and developmental conditions (∼63%) ^2,5^. Specifically, de novo *NRXN1* exon 19 deletions showed high penetrance for an autism diagnosis ^2^. Most cases carry mono-allelic deletions. However, bi-allelic deletions also occur, either as homozygous (identical mutations on both alleles) or compound heterozygous deletions (different mutations on each allele), and are often associated with more severe phenotypes ^2,6,7^. Mono-allelic mutations exhibit variable penetrance and may occur in typical individuals, or in individuals with autism, ID, schizophrenia, or other neurodevelopmental conditions ^6^. This variability is thought to arise from gene-gene and gene-environment interactions that modulate clinical outcome ^6^. In contrast, penetrance of bi-allelic NRXN1 deletions is more consistent and associated with severe ID, severe epileptic encephalopathy, Pitt Hopkins syndrome and autistic traits ^7,8^. Together, these observations suggest that NRXN1 is dosage-sensitive and that bi-allelic disruption may reveal developmental functions not apparent in mono-allelic states ^9^.

Functional genomic studies of *NRXN1* loss-of-function have been associated with impaired calcium-dependent neurotransmitter release ^10,11^, reduced expression of postsynaptic neurotransmitter receptors ^12,13^, and disruptions in synaptic plasticity ^14^. Loss of NRXN1-mediated interactions with neuroligins can also alter neuronal arborisation and modify axonal and dendritic growth patterns ^14,15^. Human induced pluripotent stem cells (iPSCs) have been widely used to model the role of NRXN1 in human neurodevelopment and its contributions to the aetiology of autism and schizophrenia ^16,17^. Most studies have focused on heterozygous mutations, either derived from affected individuals or introduced through gene editing ^16^. Engineered exon 19 mutations, for example, impair synaptic transmission by disrupting presynaptic calcium signalling and reducing neurotransmitter release in excitatory neurons ^18,19^. Of note, these studies report minimal impact on neuronal differentiation or synapse formation ^9,16^. To date, few studies have examined the consequences of bi-allelic *NRXN1* mutations or directly compared mono- and bi-allelic loss-of-function in human neurons.

To better understand the impact of bi-allelic *NRXN1* mutations and *NRXN1* gene dosage on neurodevelopment, we generated isogenic mono-allelic and bi-allelic mutations in *NRXN1* exon 19 using an iPSC line containing the transcription factor NGN2 under the control of the OPTi-OX (optimised inducible overexpression) system^20^. Isogenic mutations were introduced using a non-homologous end joining CRISPR-Cas9 approach, resulting in mono-allelic or compound bi-allelic frameshift in exon 19. Wildtype and mutant OPTi-OX were differentiated into glutamatergic neurons (herein referred to as ioGlutamatergic neurons) through induced expression of NGN2 ^20^ and molecular, cellular and functional assays were performed to characterise the resulting neurons. Using previously published work ^18,19^, we benchmarked phenotypes observed in our mono-allelic mutant lines against established findings and assessed whether bi-allelic mutations resulted in more severe cellular consequences. We find that our mono-allelic mutant lines displayed consistent molecular and cellular phenotypes with previous reports ^18,19^ and that isogenic cell lines carrying bi-allelic compound heterozygous deletions exhibited more severe phenotypes. These findings suggest that *NRXN1* mutations show gene dose sensitivity, with bi-allelic disruption in exon 19 resulting in more pronounced molecular and cellular phenotypes.

## Results

### Characterisation of *NRXN1* isoform expression in wild type and mutant ioGlutamatergic neurons

To understand the expression pattern of *NRXN1* and related genes in developing neurons, we generated glutamatergic neurons (ioGlutamatergic neurons) and first assessed the expression of two major *NRXN1* isoforms: *NRXN1α* and *NRXN1β*, at time points: 0 (iPSCs stage), 7, 14 and 21 days of differentiation **(Figure 1A)**. *NRXN1α* expression increased gradually from iPSC stage until day 21 while the *NRXN1β* isoform rose sharply at day 7 and remained at a similar level through day 21 (**Figure 1B**; mixed-effects model with Geisser–Greenhouse correction: Time – F(1.164, 6.983) = 4.04, p = 0.08; Isoform – F(2, 6) = 0.15, p = 0.86; Time × Isoform – F(6, 18) = 1.39, p = 0.27). Post hoc comparisons showed a trend toward increased *NRXN1α* expression at day 21 compared to day 0 (p = 0.10) and between day 21 and day 7 (p = 0.05), while *NRXN1β* expression remained stable after an early rise (all p > 0.1). These temporal expression patterns are consistent with those observed in fetal tissue where *NRXN1β* displays higher expression relative to *NRXN1α* ^21^. We next examined the expression profile of *NRXN2*, *NRXN3* as well as *NLGN1*, *NLGN2*, *NLGN3* and *CASK* **(Figure 1C)**. All genes showed a progressive increase in expression as ioGlutamatergic neurons matured, with NLGN2 and NLGN3 exhibiting the greatest increased relative to the iPSC stage (**Figure 1C**; one-way repeated-measures ANOVA with Geisser–Greenhouse correction: *NLGN2* – F(3, 6) = 8.43, *p* < 0.05; *NLGN3* – F(1.196, 2.392) = 15.26, *p* = 0.044; all others *p* > 0.1; Dunnett’s post hoc vs day 0: *NLGN2* day 14 *p* < 0.05, day 21 *p* < 0.001; *NLGN3* day 14–21 *p* < 0.05). Conversely, *CASK* expression remained stable across all time points following differentiation (F(1.175, 2.350) = 4.10, *p* = 0.16). These data demonstrate that ioGlutamatergic neurons progressively upregulate multiple NRXN1-related synaptic adhesion molecules during differentiation, supporting their suitability as a model for investigating NRXN1 and its molecular phenotypes.

**Figure 1:**
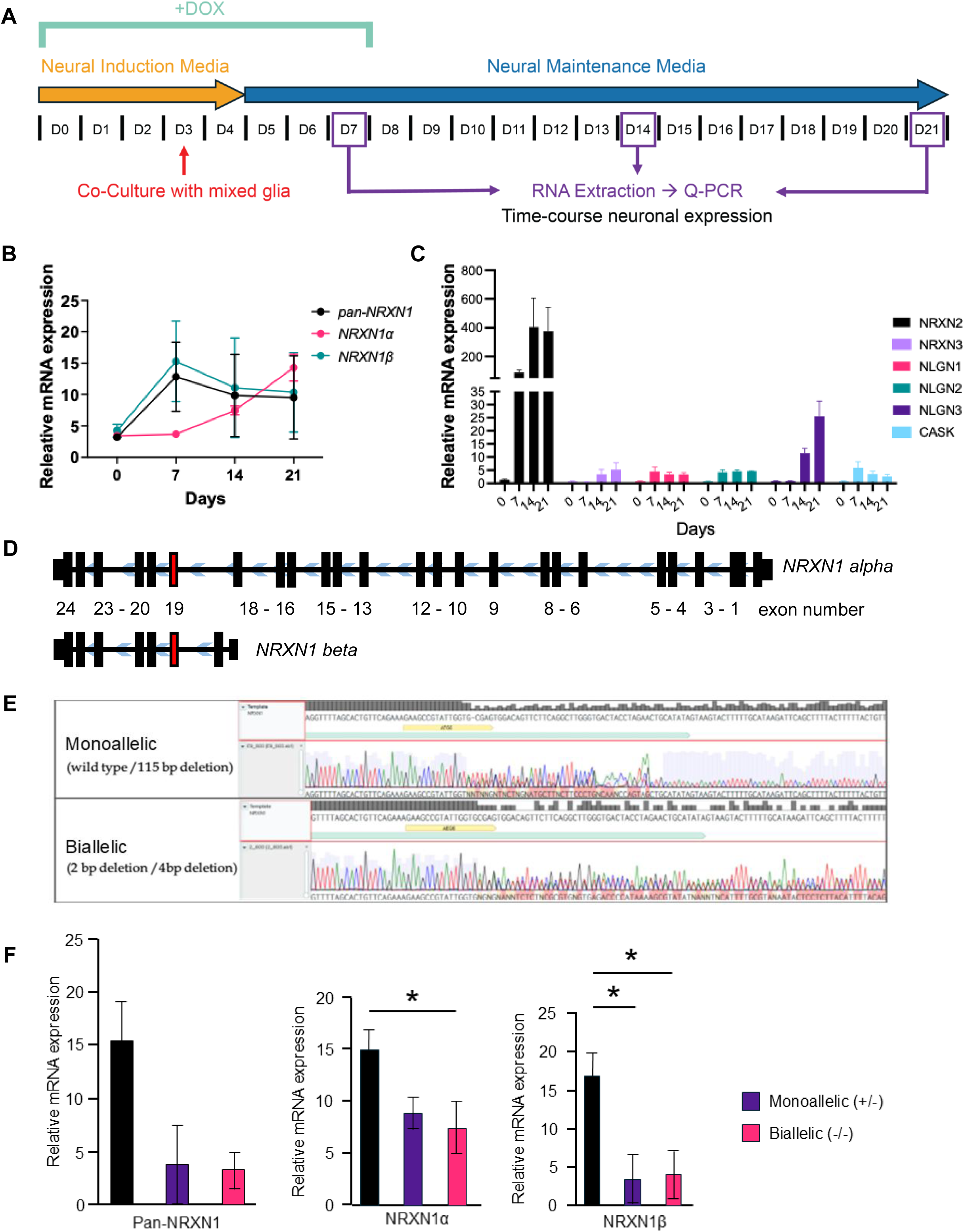
Generation of mono-allelic and bi-allelic NRXN1 mutant iPSC lines, and differentiation into iNeurons. (A) Differentiation timeline of ioGlutamatergic neurons. Characterisation of expression of NRXN1 and related genes was performed at day7, day14 and day21 of differentiation. (B) NRXN1α expression increased gradually from iPSC stage until day 21 while the NRXN1β isoform rose sharply at day 7 and remained at a similar level through day 21. (C) Expression profile using Q-PCR of NRXN2, NRXN3 as well as NLGN1, NLGN2, NLGN3 and CASK. (D) We targeted exon 19 to achieve NRXN1 loss of function as it deletes both major isoforms NRXN1α and NRXN1β. (E) At day21 of differentiation, both NRXN1α and NRXN1β and pan-NRXN1 was reduced.

Mono-allelic *NRXN1* mutations are the most frequently observed *NRXN1* gene mutations and are thus most commonly studied ^10,18,19,22,23^. Bi-allelic mutations have also been reported ^6,16^ although their effect on synapse activity is not well understood. Exon 19 is the most commonly targeted genomic region to achieve *NRXN1* loss of function as it deletes both major isoforms *NRXN1α* and *NRXN1β* ^18,19^ **(Figure 1D)**. We used a non-homologous end joining CRISPR-Cas9 approach to introduce indel mutations in OPTi-OX iPSCs within *NRXN1* exon 19. A gDNA targeting *NRXN1* exon19 (Chr2: 50,236,786-50,236,973) was electroporated into OPTi-OX iPSCs and the resulting clones genotyped (**Figure 1D** **& E**). We were successful in generating a mono-allelic deletion (wild type / 115 bp deletion) and a bi-allelic, specifically a compound heterozygous deletion (2 bp deletion / 4bp deletion) in our iPSCs **(Figure 1E)**. NGN2 OPTi-OX iPSCs underwent doxycycline-induced forward programming into NGN2-expressing neurons with glutamatergic properties (ioGlutamatergic neurons). Assessment of pluripotency, NGN2 and neuronal marker expression in isogenic control and mutant NRXN1-derived ioGlutamatergic neurons revealed that all cell lines displayed similar patterns of expression for these genes (**Supplemental Figure 1**). This indicated that the introduction of indels in the *NRXN1* gene did not alter the differentiation capacity of isogenic clones.

Next, we assessed the impact of introduced indels on *NRXN1α* and *NRXN1β* isoform expression. Across all lines *pan-NRXN1* levels were reduced compared to unedited control cells at day 21. Mutant *NRXN1*-derived ioGlutamatergic neurons also displayed significantly reduced levels of both *NRXN1α* and *NRXN1β* isoforms at day 21 (**Figure 1F**; one-way ANOVA: <u>pan-NRXN1 – F(4,10) = 2.9, P=0.075;</u> *NRXN1α* – F(4,10) =3.72, p<0.05, Dunnett’s post hoc, *p<0.05; *NRXN1β* - F(4,10) =5.12, p<0.05, Dunnett’s post hoc, *p<0.05, **p<0.01). Together these data indicate that both mono-and bi-allelic mutations in NRXN1 exon 19 impact major isoform expression, without altering the overall development of ioGlutamatergic neurons.

### Transcriptomic profiling of mutant *NRXN1* ioGlutamatergic neurons reveals disrupted neurodevelopmental pathways

Although NRXN1 is primarily known as a cell adhesion molecule which regulates synaptic transmission, a number of independent studies have shown that loss of *NRXN1* may also affect gene pathways associated with brain development and patterning, development of axons, and regulation of neuronal projection ^18,24–26^. Therefore, to gain an insight into any potential molecular disturbances caused by the introduction of exon19 *NRXN1* mutations, we performed transcriptomic analysis on day 21 ioGlutamatergic neurons generated from isogenic control, mono-allelic or bi-allelic mutant lines. Both mutant-*NRXN1* lines displayed significant alterations in their transcriptomic landscape **(Figure 2A, 2B)**. We found that neurons generated from the bi-allelic mutant line yielded greater differential expression (Number of DE genes: 1940; up=1714, down=226) compared to neurons from the mono-allelic mutant line (Number of DE genes: 247;up=69, down=178). This suggests a greater impact of the bi-allelic deletion on the transcriptional machinery compared to the mono-allelic mutations, despite having similar effects on *NRXN1* isoform expression.

**Figure 2:**
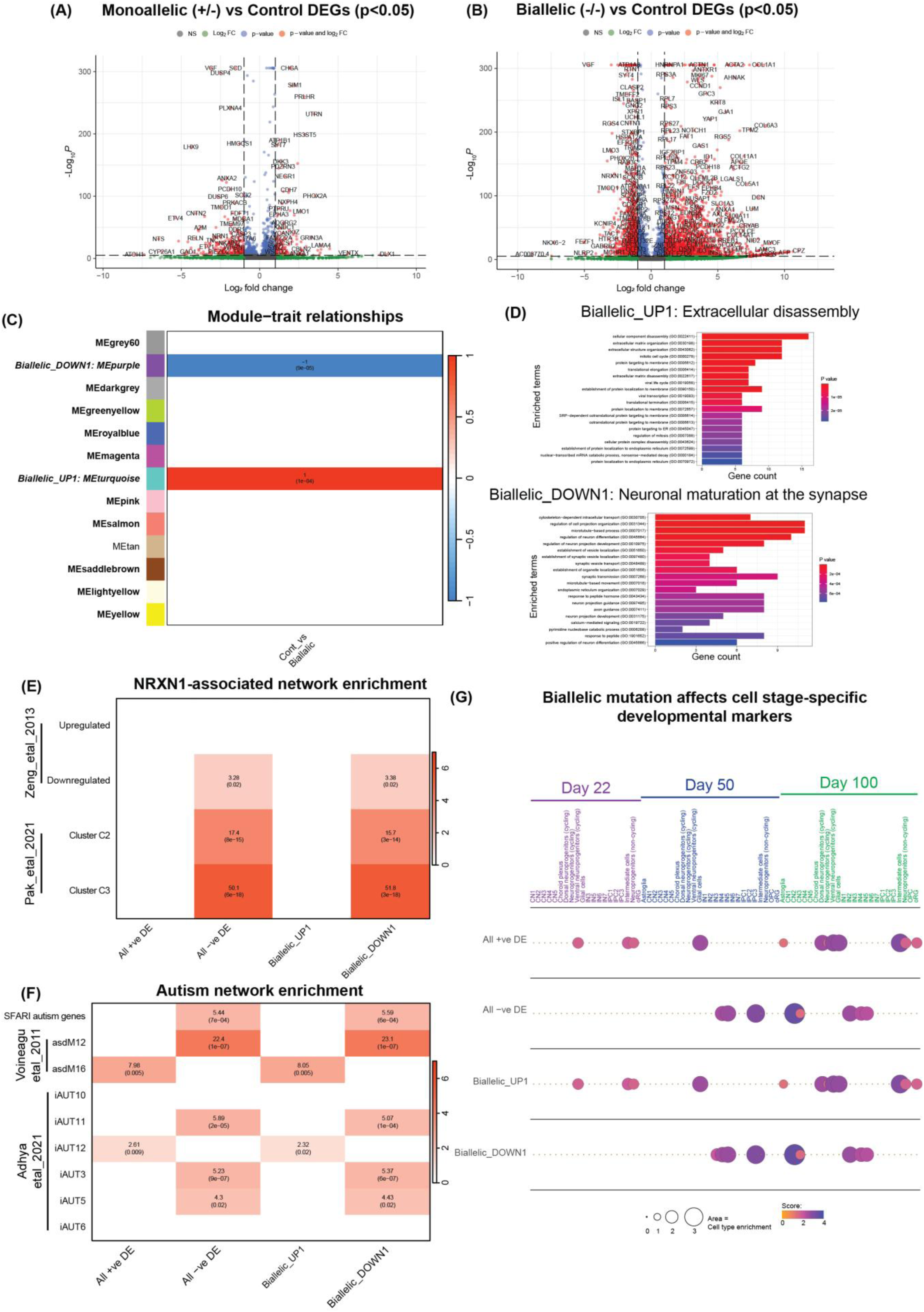
Transcriptomic profiling of mutant NRXN1 ioGlutamatergic neurons. (A) A monoallelic mutation results in differential expression of total 247 genes (up=69, down=178) as shown in this volcano plot. (B) A biallelic mutation results in differential expression of total 1914 gene (up=1714, down=226). (C) WGCNA reveals one positively correlated (Biallelic_UP1) and one negatively correlated (Biallelic_DOWN1) gene network disrupted as a result of a biallelic mutation. (D) GO analysis of co-expression modules revealed Biallelic_UP1 was enriched for extracellular disassembly related functions, while Biallelic_DOWN1 was enriched for dysregulation of neuronal maturation at the synapse. (E) NRXN1-associated network enrichment with two prominent studies that have looked at both engineered and naturally occurring mutations. (F) Enrichments with autism gene networks associated with neuronal dysfunction observed both in post mortem brains and iPSC-derived neurons. (G) Biallelic mutation affects cell-stage specific markers of brain development.

We next used an unsupervised analysis approach – Weighted Gene Co-expression Network Analysis (WGCNA) – to identify altered biological processes by constructing gene networks to identify modules of highly co-expressed genes. We did not find any dysregulated gene networks in the mono-allelic mutant neurons. However, we identified two significant dysregulated gene networks in the bi-allelic mutant neurons, one positively correlated (Biallelic_UP1) and the other negatively correlated (Biallelic_DOWN1) **(Figure 2C)**. GO analysis of co-expression modules revealed Biallelic_UP1 was enriched for extracellular disassembly related functions, while Biallelic_DOWN1 was enriched for dysregulation of neuronal maturation at the synapse **(Figure 2D)**.

Published studies on *NRXN1* mutations observed significant dysregulation only in specific pathways ^18,26^ and this corroborated with our data. We quantified the overlap of dysregulated pathways found in these study with previously published gene networks using enrichment analysis. First we ran enrichments with a study which used conditional knockdown of *NRXN1* and reported dysregulation of gene pathways associated with cell adhesion and neuron differentiation, where dysregulated pathways were downregulated ^26^. Unsurprisingly, we found a significant enrichment of these genes with the overall downregulated genes in our study **(Figure 2E)**. We also found our ‘Biallelic_DOWN1’ gene network (pathways associated with neuronal maturation at the synapse) enriched for these genes **(Figure 2E)**, suggesting specific synapse-associated pathways were only being affected. Since our *NRXN1* mutations were engineered, we wanted to test if gene pathways dysregulated as a result correlated with naturally occurring *NRXN1* mutations. Pak et al (2021) found clusters of dysregulated genes in their iPSCs from *NRXN1* participants that were functionally associated with synaptic signalling and presynaptic function ^18^. From our enrichment analysis, we found the overall downregulated genes and the Biallelic_DOWN1 module to be significantly enriched in the C2 and C3 gene clusters from Pak et al 2021 **(Figure 2E)**. *NRXN1* mutations such as those reported in Pak et al 2021 are associated with schizophrenia. However, a mutation in exon 19 is most strongly associated with autism. To quantify an association with autism genes and networks, we undertook enrichments with autism gene networks associated with neuronal dysfunction observed both in post mortem brains and iPSC-derived neurons ^27,28^ **(Figure 2F)**. Overall downregulated genes and those in the Biallelic_DOWN1 module were also significantly enriched in downregulated pathways in these published studies, while in addition overall upregulated genes and Biallelic_UP1 were enriched in upregulated pathways in these studies. Enrichment analysis also revealed enriched SFARI autism likelihood genes in our downregulated genes strongly suggesting attenuation of gene activity associated with autism **(Figure 2F)**.

In a study using *NRXN1* mutant neural organoids, the authors revealed detailed single cell transcriptomic signatures of brain development associated with NRXN1 loss of function ^25^. To understand the significance of a biallelic mutation on equivalent stages of brain development we ran stage-specific correlation of our gene networks with those identified in the neural organoid study. The overall downregulated genes and our Biallelic_DOWN1 gene network were most enriched for day50 (equivalent to early second trimester; GW14-16) and day100 (early third trimester; GW28-30) developmental stages suggesting that it is during these stages of brain development that neurons are likely to be vulnerable to the downstream effects of *NRXN1* mutations **(Figure 2G)**. The enrichment of the upregulated genes were less well-defined with developmental stages with poorer representations confirming a greater loss of function effect on brain development **(Figure 2G)**.

Overall, mono-allelic NRXN1 mutations produced limited transcriptional changes, whereas biallelic disruption was associated with extensive differential expression and two dysregulated gene modules enriched for synaptic maturation–related processes. Enrichment analyses showed overlap with prior NRXN1 perturbation datasets, with organoid stage–specific signatures corresponding to GW14–16 and GW28–30, and an overrepresentation of autism-associated gene sets.

### Bi-allelic mutations impact synaptogenesis in ioGlutamatergic neurons with greater severity

Previous studies examining the impact of *NRXN1* haploinsufficiency in exon 19 using human neurons revealed no effect on overall neuronal or synaptic morphology ^18,19^. Interestingly, our transcriptomic analysis suggests disruption of genes associated with cytoskeletal function and synaptic function in mutant neurons carrying bi-allelic mutations. Thus, we examined both neuronal and synaptic morphology in both mono-and bi-allelic mutant neurons. Consistent with previous studies, day 6 ioGlutamatergic neurons generated from both mono- and bi-allelic mutant lines did not display any differences in MAP2-positive neurite number, length or branch point (**Supplemental Figure 2**). We next assessed the synaptic architecture of ioGlutamatergic neurons generated from control, mono- or bi-allelic *NRXN1* mutant lines grown until day 21. Synapses were identified by immunostaining for the pre-synaptic protein, SYNAPSIN1 and post-synaptic protein, HOMER1 (**Figure 3A**). Synaptic proteins were imaged using an Airyscan super-resolution confocal technique which offers a lateral resolution of ∼150nm, which is greater than traditional confocal microscopy. Surprisingly, this revealed a significant increase in pre-synaptic SYNAPSIN1 density in neurons generated from mono-allelic mutant lines compared to control ioGlutamatergic neurons (**Figure 3B**). Conversely, ioGlutamatergic neurons from bi-allelic mutant neurons displayed reduced SYNAPSIN1 density compared to neurons generated from control or mono-allelic mutant lines. Furthermore, SYNAPSIN1 puncta were significantly larger in neurons generated from bi-allelic mutant lines. Examination of the post-synaptic HOMER1 puncta revealed that both mono- and bi-allelic mutant lines had increased density of HOMER1. No effect on puncta size was observed. Together, these data indicate that bi-allelic mutation had a more pronounced impact on synaptogenesis compared to mono-allelic mutation.

**Figure 3:**
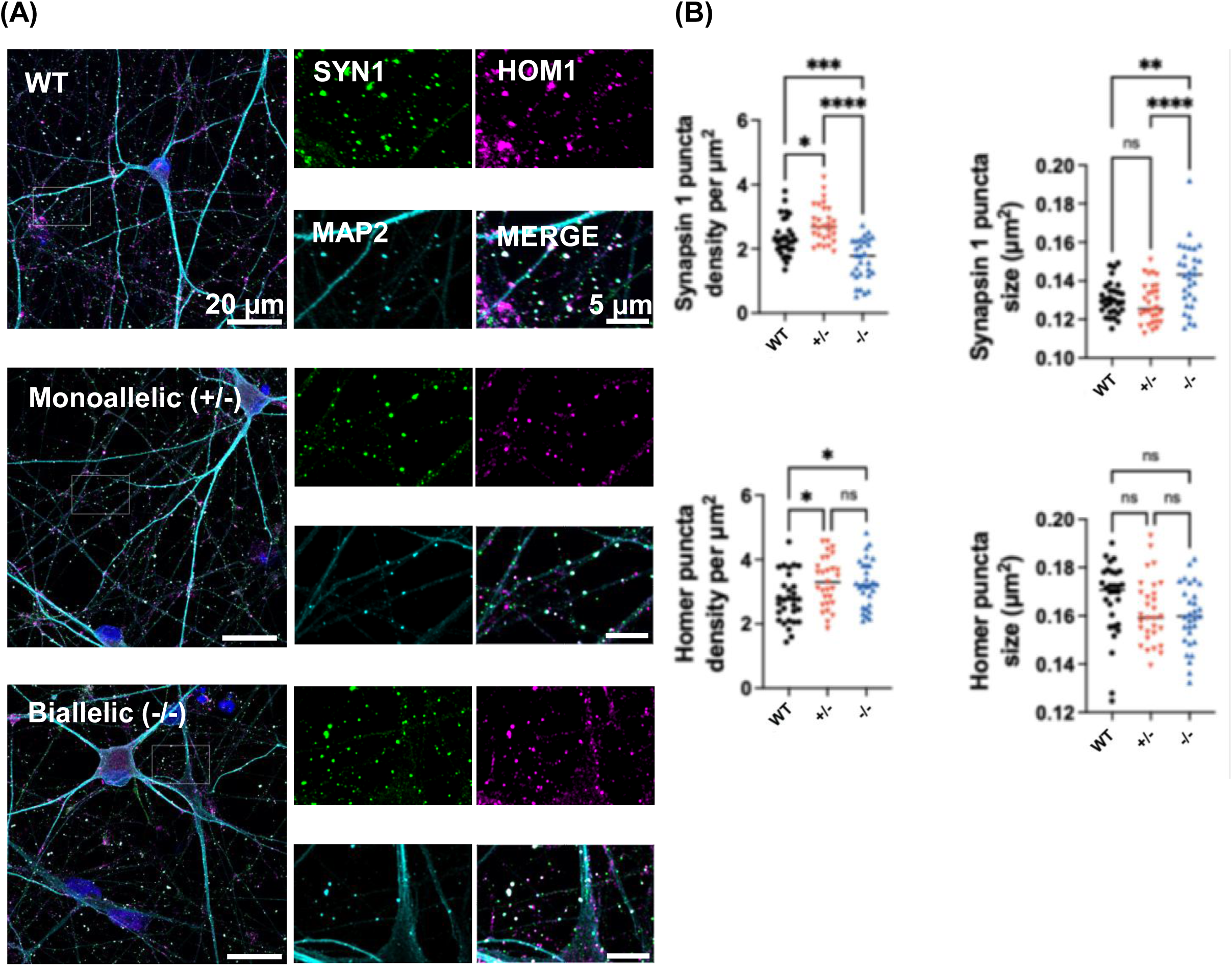
Synaptogenesis in monoallelic and biallelic mutant ioGlutamatergic neurons. (A) Synaptic proteins were imaged using an Airyscan super-resolution confocal technique which offers a lateral resolution of ∼150nm. (B) Effect of monoallelic and biallelic mutations on puncta density and size using both pre-synaptic (SYNAPSIN1) and post-synaptic markers (HOMER1).

### Bi-allelic *NRXN1* mutant lines display altered neuronal network activity

To determine whether mono- or bi-allelic *NRXN1* mutations differentially impact the development of neuronal network activity, we assessed spontaneous electrophysiological activity using multi-electrode arrays (MEAs) from day 7 to day 49 of differentiation (**Figure 4A–E**). Mean firing rate (MFR), a general measure of spontaneous neuronal activity, increased significantly over time across all genotypes (two-way repeated-measures ANOVA: main effect of time, F(1.113, 67.86) = 18.99, p < 0.0001; Figure 4A). A significant main effect of genotype was also observed, with NRXN1 mutant neurons exhibiting higher overall MFR compared to controls (F(2, 61) = 5.148, p = 0.0086). Although the time × genotype interaction did not reach statistical significance (F(2.225, 67.86) = 2.974, p = 0.0524), post hoc analysis revealed elevated MFR in mono-allelic mutant neurons from day 14 onwards and in bi-allelic mutant neurons from day 28 onwards, with no significant differences detected between the two mutant genotypes at individual time points. We next examined bursting behaviour, which reflects mesoscale organisation of neuronal firing. Burst frequency was significantly increased in both mono-allelic and bi-allelic NRXN1 mutant neurons compared to controls at later developmental stages, including day 35 and day 49 (**Figure 4B**). No significant differences were detected between mono-allelic and bi-allelic mutants though, indicating a stable genotype-dependent alteration in network bursting. Burst frequency was not significantly affected by mono- or bi-allelic genotype, developmental time, or their interaction (mixed-effects model; all p > 0.05; **Figure 4B**), suggesting that *NRXN1* mutations preferentially affect coordinated network bursting rather than intrinsic burst generation. Finally, we assessed network synchronisation, a higher-order measure of coordinated activity across electrodes. The synchrony index increased over time across all genotypes (two-way repeated-measures ANOVA: main effect of time, F(1.211, 75.10) = 20.71, p < 0.0001) and differed significantly by genotype (F(2, 62) = 5.483, p = 0.0064; **Figure 4B**). Importantly, a significant time × genotype interaction was observed (F(2.423, 75.10) = 5.103, p = 0.0053), indicating divergent developmental trajectories of network synchronisation. Post hoc analysis revealed increased synchrony in mono-allelic mutant neurons compared to controls from day 35 onwards, and in bi-allelic mutant neurons from day 42 onwards, while no significant differences were detected between mono- and bi-allelic mutants at individual time points.

**Figure 4:**
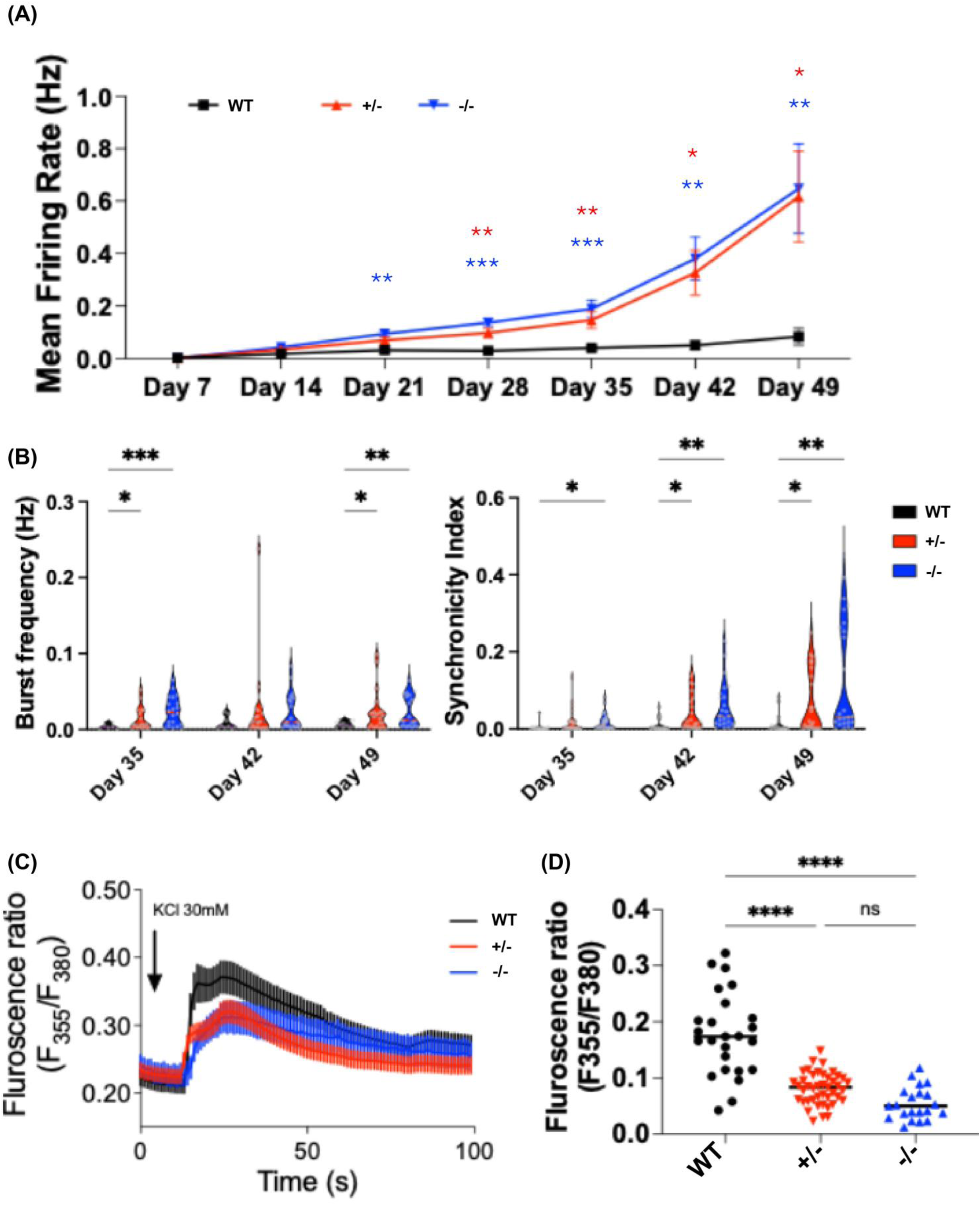
Neuronal activity in mutant NRXN1 ioGlutamatergic neurons. (A) Mean firing rate in mutant ioGlutamatergic neurons. (B) Burst frequency and synchronicity index in ioGlutamatergic neurons. (C) Changes in peak amplitude of fluorescence ratio (F355/F380) following exposure to KCl (time course). (D) Differences in peak amplitude of fluorescence ratio between control, monoallelic and biallelic mutant neurons.

Together, these data demonstrate that NRXN1 mutations disrupt neuronal network function across multiple hierarchical levels of activity. While both mono- and bi-allelic mutations increase overall excitability and bursting, bi-allelic disruption is associated with a more pronounced and divergent trajectory of network synchronisation, consistent with gene-dosage–dependent effects on higher-order network organisation.

### Mono-allelic and bi-allelic mutations both reduce peak excitability but do not impair depolarisation

Previous reports indicate that NRXN1 mutations exert distinct effects on different aspects of neuronal signalling, with increased network-level activity observed alongside impaired synaptic transmission and evoked responses ^17,29^ ^19^. To assess depolarisation in our NRXN1 mutant neurons, we measured intracellular calcium signalling in response to KCl stimulation, as this is an indicator of neuronal excitability and calcium release efficiency during neuronal depolarisation. We recorded changes in peak amplitude of fluorescence ratio (F355/F380) following exposure to KCl. Compared to our control neurons, we observed significant reduction in peak amplitudes in both mono-allelic and bi-allelic mutant neurons (Peak amplitude: WT, 0.177 ± 0.015; mono-allelic, 0.08 ± 0.004; bi-allelic, 0.055 ± 0.007) **(Figure 4C and D)**. This reduction suggests impaired efficiency of depolarisation-evoked calcium influx, rather than a global reduction in neuronal activity. However, despite reduced peak amplitudes, the overall calcium response profiles following 30 mM KCl stimulation were preserved and followed similar temporal dynamics across control and mutant neurons, indicating that depolarisation responses were maintained but altered in magnitude (**Figure 4C**). Thus, NRXN1 mutant neurons retain the capacity to respond to excitatory stimuli, but exhibit attenuated peak calcium entry.

Taken together, these findings indicate a dissociation between increased network-level excitability and reduced peak depolarisation responses at the cellular level in NRXN1 mutant neurons. This interpretation is consistent with previous work showing that NRXN1 mutations reduce the amplitude of initial evoked responses without abolishing subsequent depolarisation events ^19^. Such a mechanism could explain why neuronal excitability was not reduced in NRXN1 mutant individuals ^29^, and why we observed increased excitability in our neuronal cultures.

## Discussion

Mono-allelic *NRXN1* mutations are associated with marked variability in clinical presentation, whereas bi-allelic mutations are consistently linked to severe neurodevelopmental phenotypes ^6^ ^7,8^. Despite this, the majority of functional studies, including nearly all human iPSC-based models, have focused on mono-allelic *NRXN1* mutations ^16^. This raises the possibility that certain *NRXN1*-dependent phenotypes may be attenuated or masked by the presence of a functional allele and therefore remain undetected in heterozygous systems. In this study, we addressed this gap by directly comparing isogenic mono-allelic and bi-allelic *NRXN1* exon 19 loss-of-function mutations in human glutamatergic neurons generated using the OPTi-OX system.

Previous iPSC studies of mono-allelic *NRXN1* mutations have consistently reported subtle and context-dependent functional effects. Conditional exon 19 disruption in gene-edited iPSCs revealed transient impairments in evoked neurotransmitter release, characterised by reduced response amplitude and altered release probability following stimulation, without detectable effects on neuronal differentiation or synaptic structure ^19^. Participant-derived iPSC neurons carrying *NRXN1* mutations similarly showed changes in firing dynamics, but no consistent alterations in synapse density or dendritic morphology ^18^. In parallel, studies using neurons derived from *NRXN1* haploinsufficient individuals reported increased neuronal excitability without changes in action potential threshold, alongside modest transcriptional perturbations enriched for synaptic pathways ^29^.

Our findings in mono-allelic *NRXN1* mutant neurons are broadly consistent with this literature. We found modest gene expression alterations, which did not enrich in specific pathways or gene networks. This is similar to gene expression alterations from participant-derived iPSC-neurons ^18,25,29^. Moreover, we observed reduction of peak excitability, similar to reduction in amplitude of action potential seen in previous studies, but we did not find evidence of SYNAPSIN1 disruption at the pre-synapse which was also noted in previous studies ^19^. However, we did observe alterations in network properties and in response to depolarising stimuli. Taken together, our data is consistent with previous reports ^18,19,25^ that mono-allelic mutations produce modest effects on molecular and cellular phenotypes, but with robust impacts on functional properties. This dissociation between modest molecular disruption and pronounced functional alterations suggests that *NRXN1* haploinsufficiency primarily affects synaptic efficiency and network dynamics rather than gross synaptic architecture or transcriptional programmes.

There is not much known about the effects of a bi-allelic *NRXN1* mutation at a molecular, cellular or functional level. We find that bi-allelic mutations produced phenotypes that overlapped with, extended beyond, or were not detected in mono-allelic mutant neurons. First, we found a very high number of differentially expressed genes in bi-allelic mutant neurons – greater than seven times compared to mono-allelic mutant neurons – and identified novel gene co-expression pathways dysregulated associated with bi-allelic *NRXN1* mutations. Further, we observed alterations in the expression patterns of both HOMER1 and SYNAPSIN1 synaptic proteins, which were less pronounced in neurons with mono-allelic mutations. Interestingly, despite these differences at the molecular and cellular level, both mono-and bi-allelic mutation carrying neurons display similar alteration in neuronal network properties and in response to depolarising stimuli. Collectively, in contrast to mono-allelic mutations, bi-allelic mutation of NRXN1 not only affects synaptic efficiency and network dynamics, but also synaptic architecture, potentially driven by alterations in transcriptional programmes.

Taken together, our findings support previous studies indicating that mono-allelic mutations in *NRXN1* have modest effects at molecular levels. We further go on to provide evidence that compound bi-allelic mutations have more pronounced impacts on molecular and cellular phenotypes compared to mono-allelic mutations. these findings support a model whereby *NRXN1* mutations have a gene-dose impact in developing neurons.

## METHODS

### Generation of isogenic NRXN1 mutant iPSC lines

CRISPR–Cas9 genome editing was used to introduce indel mutations into *NRXN1* exon 19 in a human iPSC line carrying an inducible NGN2 transgene under the OPTi-OX system (hereafter NGN2-OPTi-OX iPSCs), which has been previously established and characterised ^20^. Targeting exon 19 disrupts both *NRXN1α* and *NRXN1β* isoforms.

A single guide RNA (gRNA; sequence: AGAAGCCGTATTGGTGCGAG) was designed to target the shared exon. gRNA synthesis and CRISPR–Cas9 nucleofection were performed by Aelian Biotechnology (Vienna, Austria) using a non-homologous end-joining strategy. Edited clones were isolated and genotyped to identify mono-allelic and compound bi-allelic frameshift mutations. Unedited parental NGN2-OPTi-OX iPSCs served as isogenic controls.

### Maintenance and neuronal differentiation of NGN2-OPTi-OX iPSCs

NGN2-OPTi-OX iPSCs were maintained as colonies in StemFlex medium under standard conditions ^30^. Neuronal differentiation was initiated (day 0) by doxycycline-induced NGN2 expression in neural induction medium. Induction medium was refreshed daily on days 0–2. On day 3, induced neurons were dissociated using Accutase, resuspended in neural maintenance medium, and counted. For most assays, neurons were co-cultured with primary rat mixed glial cells at a 1:1 neuron:glia ratio on poly-D-lysine (PDL)/Geltrex-coated substrates. Cultures were maintained in neural maintenance medium supplemented with doxycycline until day 7, after which doxycycline was withdrawn and half-media changes were performed every 2–3 days.

### Primary mixed glial cultures

Primary mixed glial cultures were prepared from P0–P2 Sprague–Dawley rat pups following established protocols with minor modifications. All procedures were conducted in accordance with UK Home Office Schedule 1 regulations. Cortices were dissected under sterile conditions and dissociated tissue was cultured for 10 days in DMEM (high glucose) supplemented with 10% fetal bovine serum and 1% penicillin–streptomycin, maintained at 37 °C, 7% CO_₂_.

### Quantitative PCR (qPCR)

Total RNA was extracted from ioGlutamatergic neurons using the Qiagen RNeasy Mini Kit. cDNA was synthesised from 1 µg RNA using the Maxima First Strand cDNA Synthesis Kit according to the manufacturer’s instructions. qPCR reactions were performed using gene-specific primers and normalised to housekeeping genes. Relative expression was calculated using the ΔΔCt method.

### RNA sequencing and transcriptomic analysis

RNA sequencing was performed on day 21 ioGlutamatergic neurons derived from control, mono-allelic, and bi-allelic NRXN1 mutant lines. Library preparation was carried out by the Wellcome–MRC Cambridge Stem Cell Institute NGS Facility using poly(A) mRNA enrichment (NEBNext Poly(A) Magnetic Isolation Module) followed by the NEBNext Ultra II Directional RNA Library Prep Kit. Library quality was assessed using an Agilent 4200 TapeStation.

Sequencing was performed on an Illumina NovaSeq 6000 platform to a depth of approximately 200–250 million reads per sample. FASTQ files were aligned to the human reference genome using STAR, and gene-level counts were generated using HTSeq. Differential expression analysis was conducted using DESeq2. Gene co-expression network analysis was performed using Weighted Gene Co-expression Network Analysis (WGCNA) ^28^ to identify modules associated with *NRXN1* genotype.

### Synaptic immunostaining and Airyscan imaging

For synaptic analysis, neurons were plated at 60 × 10^³^ cells/well on PDL/Geltrex-coated coverslips in 24-well plates and maintained until day 21. Cells were fixed with 4% paraformaldehyde (15 min, RT), permeabilised with 0.1% Triton X-100, and blocked in 10% goat serum. Primary antibodies against Synapsin-1, Homer-1, and MAP2 were applied overnight at 4 °C, followed by Alexa Fluor-conjugated secondary antibodies. Nuclei were counterstained with DAPI. Coverslips were mounted and imaged using a Zeiss LSM 980 Airyscan2 confocal microscope with a 63× oil objective (NA 1.4). Synaptic puncta were quantified using MetaMorph software. Images were thresholded based on mean + 2×SD intensity and analysed using Integrated Morphometry Analysis to quantify puncta density and morphology. Colocalisation between pre- and post-synaptic markers was assessed using correlation analysis.

### Microelectrode array (MEA) plating and neuronal culture

At day 3 post induction, NGN2-OPTi-OX iPSCs were dissociated into single cells using Accutase and replated onto CytoView MEA 48-well plates (Axion Biosystems) pre-coated with poly-D-lysine and Geltrex. Cells were seeded at a density of 50,000 cells per well as a 10 µL droplet directly over the electrode array to promote network formation. Following attachment, neuronal maturation medium was gently added, and cultures were maintained at 37 °C and 5% CO_₂_ for the duration of the experiment, with half-media changes every 2–3 days.

### MEA recordings

Spontaneous electrophysiological activity was recorded longitudinally from day 7 to day 49 post induction using the Maestro Pro MEA system (Axion Biosystems). Recordings were acquired weekly for 20 min per well under controlled environmental conditions (37 °C, 5% CO_₂_). All recordings were performed prior to media changes to minimise variability associated with acute handling effects. For each genotype, recordings were obtained from at least three independent differentiations with eight technical replicate wells per experiment.

### MEA signal processing and parameter extraction

Raw MEA data were processed using Axion Neural Metrics Tool software. Spike detection was performed using an adaptive threshold set to 5.5 standard deviations above baseline noise. Active electrodes were defined as electrodes exhibiting ≥5 spikes per minute. Burst detection at the single-electrode level was performed using a Poisson Surprise algorithm, with a minimum surprise threshold of 3. Network bursts were identified using an inter-spike interval (ISI)–based method, requiring a minimum of 35 spikes, a maximum ISI of 100 ms, and participation of at least 25% of electrodes within a well. The following parameters were extracted for each well, consistent with established MEA analysis frameworks ^31^:

- Mean firing rate (MFR): total number of detected spikes divided by recording duration (Hz), used as a measure of overall network excitability.
- Burst metrics: including single-electrode burst rate and network burst rate, reflecting mesoscale organisation of neuronal firing.
- Synchrony index: a unitless measure (0–1) of coordinated activity across electrodes, calculated using cross-correlation within a 20 ms synchrony window.

These parameters were selected to capture hierarchical levels of network organisation, spanning single-electrode activity, population bursting, and higher-order network synchronisation, in line with previous dissociated organoid and iPSC-derived neuronal MEA studies ^31^.

### MEA statistical analysis

For longitudinal analyses, mean values were calculated per well and averaged across technical replicates within each biological replicate. Temporal effects and genotype differences were assessed using two-way repeated-measures ANOVA or mixed-effects models where appropriate, with time and genotype as factors. Post hoc comparisons were corrected for multiple testing. MEA data are presented as mean ± SEM unless otherwise stated.

### Calcium dye loading and imaging

Intracellular calcium dynamics were assessed at day 21 post induction using ratiometric calcium imaging. NGN2-OPTi-OX neurons were dissociated and plated onto poly-D-lysine/Geltrex-coated 35 mm glass-bottom µ-dishes (Ibidi) at a density of 3.5 × 10^⁵^ cells per dish and maintained in neural induction medium. Cells were loaded with 5 µM Fura-2 AM (eBioscience) for 40 min at 37 °C and 5% CO_₂_ in culture medium, followed by a 30 min equilibration period in pre-warmed calcium-free HBSS to allow complete de-esterification of the dye. Imaging was performed at room temperature using a Nikon Eclipse Ti-S microscope equipped with a QIClick™ CCD camera (QImaging).

Ratiometric fluorescence imaging was performed using alternating excitation at 355 nm and 380 nm via a Dual OptoLED light source (Cairn Research), with emission collected at 510 nm (470–550 nm bandpass). Images were acquired every 3 s using MetaFluor® Fluorescence Ratio Imaging Software (Molecular Devices). Following a 60 s baseline recording, depolarisation was induced by bath application of 30 mM KCl, and imaging continued to capture evoked calcium responses.

Fluorescence intensity ratios (F355/F380) were calculated for individual neurons, corrected for background and autofluorescence, and normalised to baseline. Peak response amplitude was quantified as the maximal change in F355/F380 following KCl application and used as a measure of depolarisation-evoked calcium entry. Temporal response profiles were also analysed to assess response kinetics and overall waveform preservation across genotypes.

For each experiment, multiple cells were analysed per dish and averaged to generate a single biological replicate. Comparisons between genotypes were performed using one-way ANOVA or Kruskal–Wallis tests depending on data distribution, with post hoc correction for multiple comparisons. Data are presented as mean ± SEM.

## ACKNOWLEDGMENTS

The authors acknowledge funding support from UK Medical Research Council, Grant Nos. MR/L021064/1 (D.P.S.), MR/Y012968/1 (D.P.S.), MR/X004112/1 (D.P.S.), MR/Y008170/1 (D.P.S.), and MR/Y012968/1 (D.P.S.). D.P.S. is also a recipient of an Independent Researcher Award from the Brain and Behavior Foundation (Grant No. 25957). SBC received funding from the Wellcome Trust 214322\Z\18\Z. For the purpose of Open Access, the author has applied a CC BY public copyright licence to any Author Accepted Manuscript version arising from this submission. SBC also received funding from the Innovative Medicines Initiative 2 Joint Undertaking under grant agreement No 777394 for the project AIMS-2-TRIALS. This Joint Undertaking receives support from the European Union’s Horizon 2020 research and innovation programme and EFPIA and AUTISM SPEAKS, Autistica, SFARI. The funders had no role in the design of the study; in the collection, analyses, or interpretation of data; in the writing of the manuscript, or in the decision to publish the results. The University of Cambridge Autism Research Centre received funding for this work from the Autism Research Trust, whose legacy work is now managed by Autism Action. SBC also received funding from SFARI, the Templeton World Charitable Fund and the MRC. We are grateful to Cambridge University Development and Alumni Relations (CUDAR) for anonymous donations. Any views expressed are those of the author(s) and not necessarily those of the funders (including IHI-JU2). All research at the Department of Psychiatry in the University of Cambridge is supported by the NIHR Cambridge Biomedical Research Centre (NIHR203312) and the NIHR Applied Research Collaboration East of England. The views expressed are those of the author(s) and not necessarily those of the NIHR or the Department of Health and Social Care.The authors thank George Chenell of the Wohl Cellular Imaging Centre (King’s College London) for technical support.

## AUTHOR CONTRIBUTIONS

Conceptualization, A.M., D.P.S., D.A., M.K., and S.B.C.; methodology, A.M., N.J.F.G., D.P.S., D.A., M.K.; Investigation, A.M., A.P., N.J.F.G., L.D., D.A.,; writing—original draft, A.M., D.P.S., D.A.; funding acquisition, D.P.S., D.A., M.K., and S.B.C.; supervision, D.P.S., M.K., S.B.C.

---

**Supplementary Figure 1:**
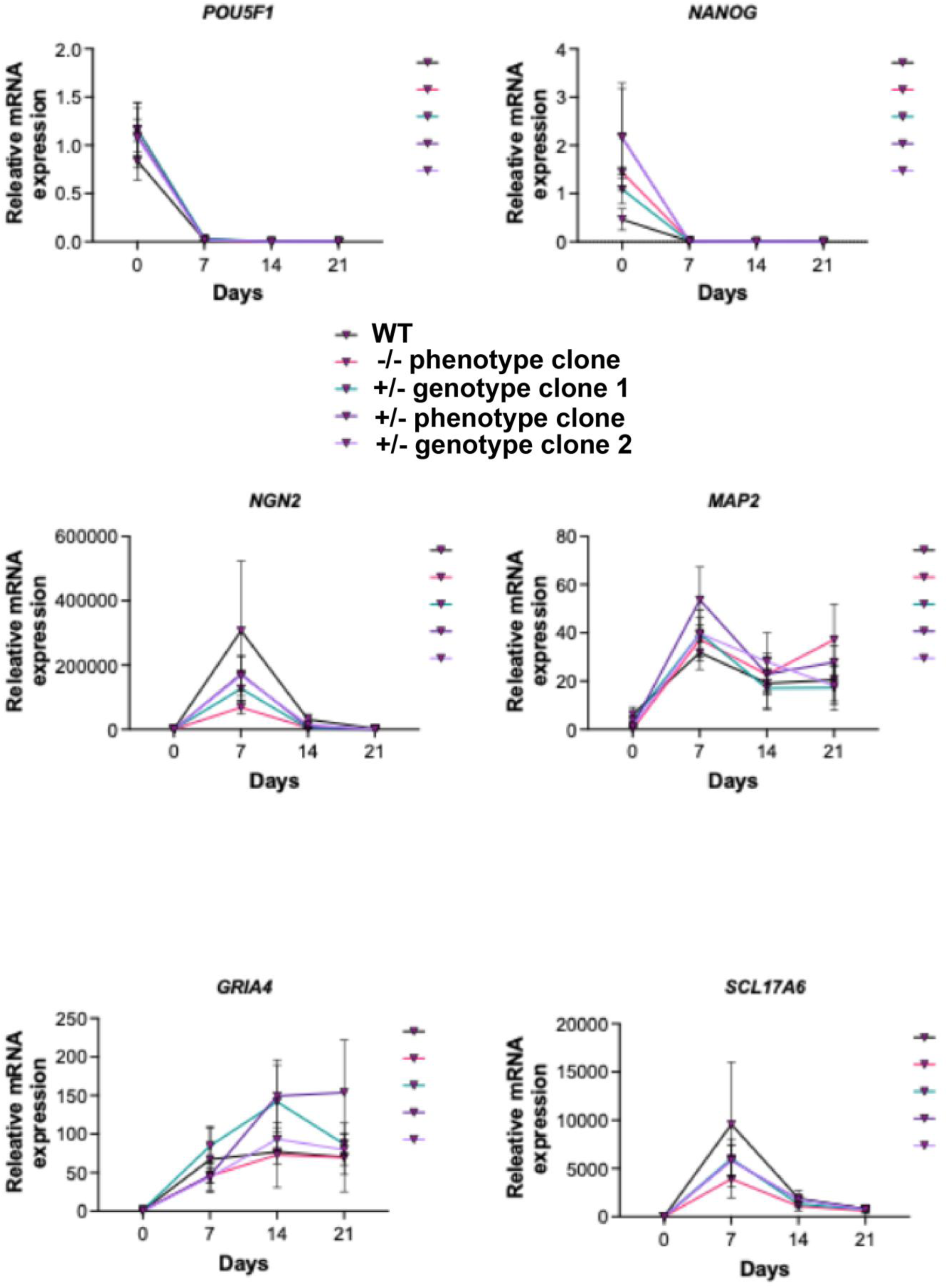
Assessment of pluripotency, NGN2 and neuronal markers expression in isogenic control and mutant NRXN1-derived ioGlutamatergic neurons.

**Supplementary Figure 2:**
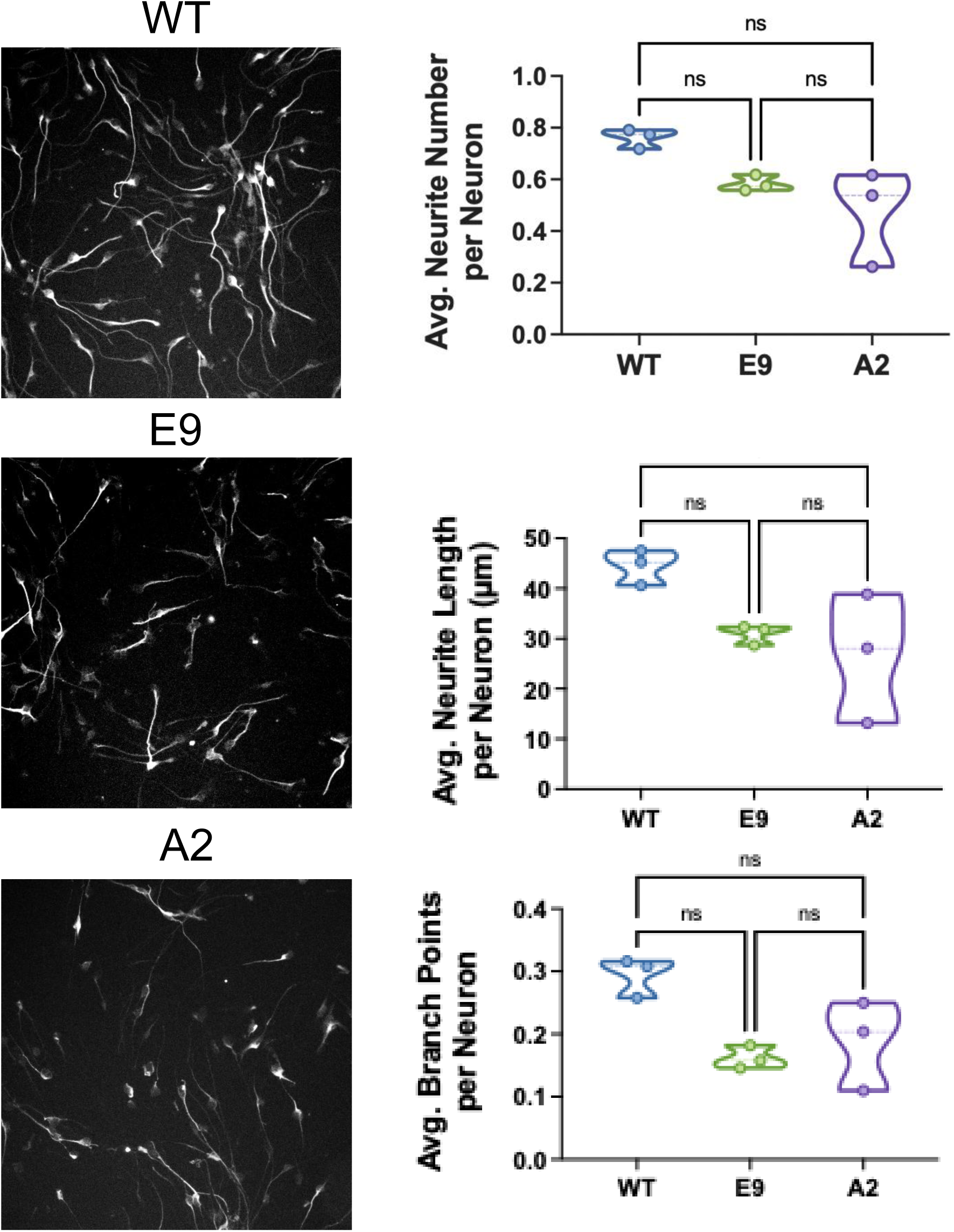
Assessment of neurite outgrowth in NRXN1-mutant neurons.

